# *cacna2d3*, a voltage-gated calcium channel subunit, functions in vertebrate habituation learning and the startle sensitivity threshold

**DOI:** 10.1101/2021.12.02.470952

**Authors:** Nicholas J. Santistevan, Jessica C. Nelson, Elelbin A. Ortiz, Andrew H. Miller, Dima Kenj Halabi, Zoë A. Sippl, Michael Granato, Yevgenya Grinblat

## Abstract

The ability to filter sensory information into relevant versus irrelevant stimuli is a fundamental, conserved property of the central nervous system and is accomplished in part through habituation learning. Synaptic plasticity that underlies habituation learning has been described at the cellular level, yet the genetic regulators of this plasticity remain poorly understood, as do circuits that mediate sensory filtering. A forward genetic screen for zebrafish genes that control habituation learning identified a mutant allele *dory*^*p177*^ that caused reduced habituation of the acoustic startle response. Whole-genome sequencing identified the *calcium voltage-gated channel auxiliary subunit alpha-2/delta-*3 (*cacna2d3*) as a candidate gene affected in *dory*^*p177*^ mutants. Behavioral characterization of larvae homozygous for two additional, independently derived mutant alleles of *cacna2d3*, together with failure of these alleles to complement *dory*^*p177*^, confirmed a critical role for *cacna2d3* in habituation learning. Notably, detailed analyses of the acoustic response in mutant larvae also revealed increased startle sensitivity to acoustic stimuli, suggesting a broader role for *cacna2d3* in controlling innate response thresholds to acoustic stimuli. Taken together, our data demonstrate a critical role for *cacna2d3* in sensory filtering, a process that is disrupted in human CNS disorders, e.g. ADHD, schizophrenia, and autism.

## Introduction

To successfully navigate their environments, animals continuously adjust their behaviors to ensure that they are appropriate for the current environment. Their nervous systems must quickly process incoming stimuli to distinguish relevant from irrelevant information, which allows for focused attention and supports higher executive functions like memory formation and behavioral regulation. Sensory filtering is mediated in part by the fundamental and conserved process of habituation (Ramaswami, 2014). Habituation, the simplest form of non-associative learning exhibited by all animals, is defined as a progressive decline in responsiveness to repeated, insignificant stimuli (Groves and Thompson, 1970), and is not due to sensory adaptation or motor fatigue (Rankin et al., 2009). The behavioral parameters and cellular mechanisms of habituation are controlled by synaptic plasticity mechanisms that alter neurotransmitter signaling to regulate a balance of excitation and inhibition (Aljure et al., 1980; Castellucci and Kandel, 1974; Marsden and Granato, 2015; Simons-Weidenmaier et al., 2006; Weber et al., 2002), but our knowledge of the critical genes that mediate habituation is incomplete.

Impairment of filtering mechanisms is a hallmark of many common neurological disorders, so much so that habituation deficits have been used as a diagnostic tool (McDiarmid et al., 2017). Habituation deficits are associated with autism spectrum disorders (ASD) (Ornitz et al., 1993; Perry et al., 2007; Takahashi et al., 2016), Fragile X syndrome (Castren et al., 2003), schizophrenia (Braff and Geyer, 1990), Huntington’s disease (Agostino et al., 1988), attention deficit hyperactivity disorder (ADHD) (Jansiewicz et al., 2004), Parkinson’s disease (Penders and Delwaide, 1971), Tourette’s syndrome (Bock and Goldberger, 1985), and migraine (Coppola et al., 2013). Dissecting the underlying genetic mechanisms that regulate sensory filtering can provide insight into the etiology of disease, identify genetic predispositions for diseases, and identify potential therapeutic targets. Importantly, understanding the genetic, cellular, and behavioral aspects of habituation is critical to understanding how normal neural circuits process sensory information.

Zebrafish can perform sensory-evoked motor behaviors that are modulated by experience by 5 days post-fertilization (dpf). Acoustic stimuli elicit one of two distinct motor responses in zebrafish: a short-latency C-bend (SLC), generally performed in response to a high-intensity stimulus and a long-latency C-bend (LLC), generally performed in response to a low-intensity stimulus (Burgess and Granato, 2007). These behaviors are driven by simple, well-characterized circuits that are accessible to visualization and genetic manipulation (Wolman and Granato, 2012). A SLC is triggered by activating one of two bilateral Mauthner hindbrain reticulospinal neurons, the command neurons of the acoustic startle response (ASR) (Korn and Faber, 2005).

The Mauthner neuron is functionally analogous to the giant neurons of the caudal pontine reticular nucleus (PnC), which receive input from the cochlear nerve and output to motor neurons in the spinal cord to drive the startle response in mammals (Gahtan et al., 2002; Lingenhohl and Friauf, 1994; Ogino et al., 2019). While the zebrafish circuitry is simpler in comparison to that of mammals, it is this simplicity that makes it a useful tool for investigating the genetic, cellular and behavior mechanisms that underlie sensory filtering.

To identify genes that are important for mediating habituation learning, we combined a genome-wide forward genetic screen (Wolman et al., 2015) with a high-throughput platform for unbiased acoustic startle analysis (Wolman et al., 2011). This approach yielded several genes required for acoustic startle habituation, including the palmitoyltransferase *Huntingtin interacting protein 14 (hip14)* (Nelson et al., 2020), the *adaptor related protein complex 2 subunit sigma 1* gene (*ap2s1*) (Jain et al., 2018), the extracellular metalloprotease *pregnancy associated plasma protein-aa (pappaa)*, and the enzyme *pyruvate carboxylase a (pcxa)* (Wolman et al., 2015). Here, we report identification of a previously uncharacterized allele named *dory* ^*p177*^ as a mutation in the *calcium voltage-gated channel auxiliary subunit alpha-2/delta-3* (*cacna2d3*) gene, predicted to result in a premature stop codon. *cacna2d3* encodes a member of the alpha2/delta subunit family of proteins in the voltage-gated calcium channel (VGCC) complex, known to be involved in synaptic transmission and neurotransmitter release (Bauer et al., 2010; Dolphin, 2013). We show that *cacna2d3* is required for vertebrate sensory filtering, with *cacna2d3* mutant zebrafish exhibiting reduced habituation and a reduced innate startle threshold to acoustic stimuli. Collectively, these data show that *cacna2d3* plays an important role in sensory filtering by controlling both the innate acoustic startle threshold and the ability of the animal to habituate to repeated, irrelevant acoustic stimuli.

## Results

### Whole-genome sequencing identifies cacna2d3 as a candidate regulator of habituation learning

Habituation is defined as a progressive decline in responsiveness to repeated, insignificant stimuli (Groves and Thompson, 1970). Habituation is the simplest form of non-associative learning found in all animals, yet its genetic regulation remains poorly understood. To identify genes involved in habituation, we combined forward genetic mutagenesis screening with a robust, unbiased behavioral assay for acoustic startle habituation (Wolman et al., 2015). *dory*^*p177*^ was among the mutant allele collection generated through this approach. To identify the likely causative mutation underlying the *dory*^*p177*^ habituation defect, whole genome sequencing (WGS) was performed followed by homozygosity mapping to identify potentially causative mutations in the *dory*^*p177*^ line. We mapped *dory*^*p177*^ to a chromosome 11 (Figure 1A) interval containing a unique mutation in the *calcium voltage-gated channel auxiliary subunit alpha-2/delta-3* gene, *cacna2d3* (Figure 1B).

**Figure 1.**
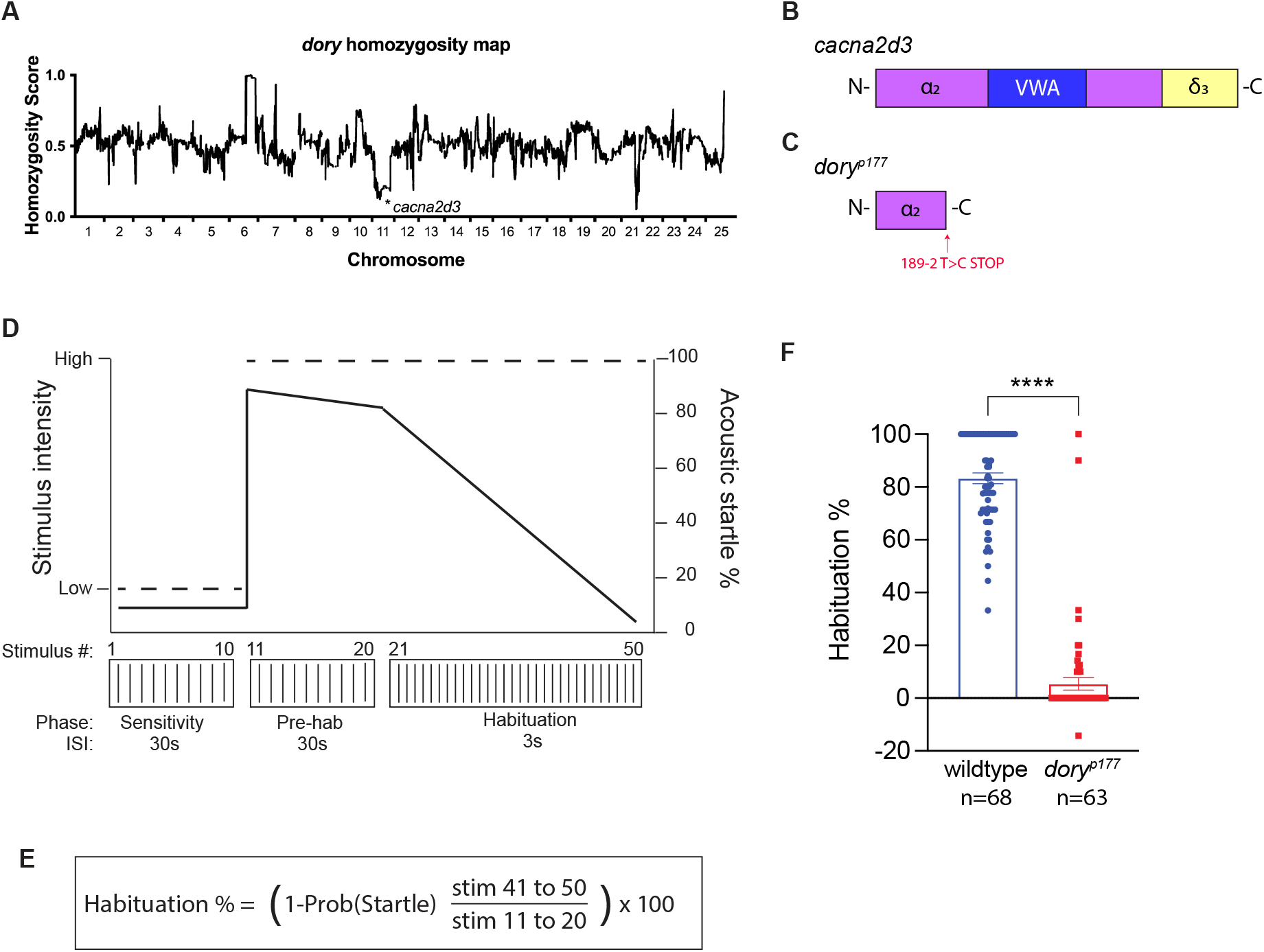
*dory*^*p177*^, linked to *cacna2d3* by homozygosity analysis, causes reduced habituation of the acoustic startle response. (A) Homozygosity plot of *dory*^*p177*^ mutants based on whole genome sequencing results. Homozygosity scores close to 1.0 indicate linkage to TL alleles while scores close to 0.0 indicate linkage to WIK alleles. Asterisk indicates the position of the identified splice-donor site mutation in *cacna2d3* on chromosome 11. (B-C) Schematics of the wildtype *cacna2d3* allele (B) and the *dory*^*p177*^ mutant allele (C). (D) Schematic representation of the acoustic startle habituation assay. Larvae were exposed to 10 “sub-threshold” low-intensity acoustic stimuli delivered at 30s interstimulus intervals (ISI) to access startle sensitivity. Next larvae were exposed to 10 high-intensity non-habituating stimuli delivered at 30s ISI to determine baseline startle responsiveness followed by 30 high intensity habituating stimuli at a 3s ISI to access habituation. The black line indicates a typical wildtype response at each phase of the assay. (E) Habituation is calculated by taking the ratio of the average frequency of startle responsiveness of an individual to stimuli 41-50 over stimuli 11-20. (F) Mean acoustic startle habituation percentage of wildtype and *dory*^*p177*^ mutants. Wildtype larvae are shown in blue and homozygous *dory*^*p177*^ mutants are shown in red. Number of larvae shown below each bar. ****p<0.0001, Mann-Whitney test versus wildtype. Error bars indicate SEM.

Cacna2d3 is a member of the alpha-2/delta subunit family of proteins, which regulate the trafficking and surface expression of voltage-gated calcium channel (VGCC) complexes; VGCCs in turn modulate synaptic transmission and function (Dolphin, 2013; Hoppa et al., 2012; Pirone et al., 2014). The primary sequence of alpha-2/delta-3 is strongly conserved across vertebrates; human CACNA2D3 shares 76.5% amino acid identity with zebrafish Cacna2d3 protein, 87.6% similarity along the length of the protein, and 89.4% amino acid identity/93.4% similarity in the Von Willebrand Factor A (VWA) functional domain (Supplemental Figure 1). The unique thymine to cytosine single base pair substitution in the *dory*^*p177*^ allele resides at the donor splice site of *cacna2d3* intron 2-3 and is predicted to cause partial retention of the intronic sequence in the mutant transcript, leading to a premature stop codon 39 bp following the mutation (Figure 1C). The predicted mutant protein lacks the VWA domain and the *δ-3* subunit that anchors Cacna2d3 to the cell membrane (Figure 2B), both critical to Cacna2d3 function in VGCCs (Dolphin, 2013).

**Figure 2.**
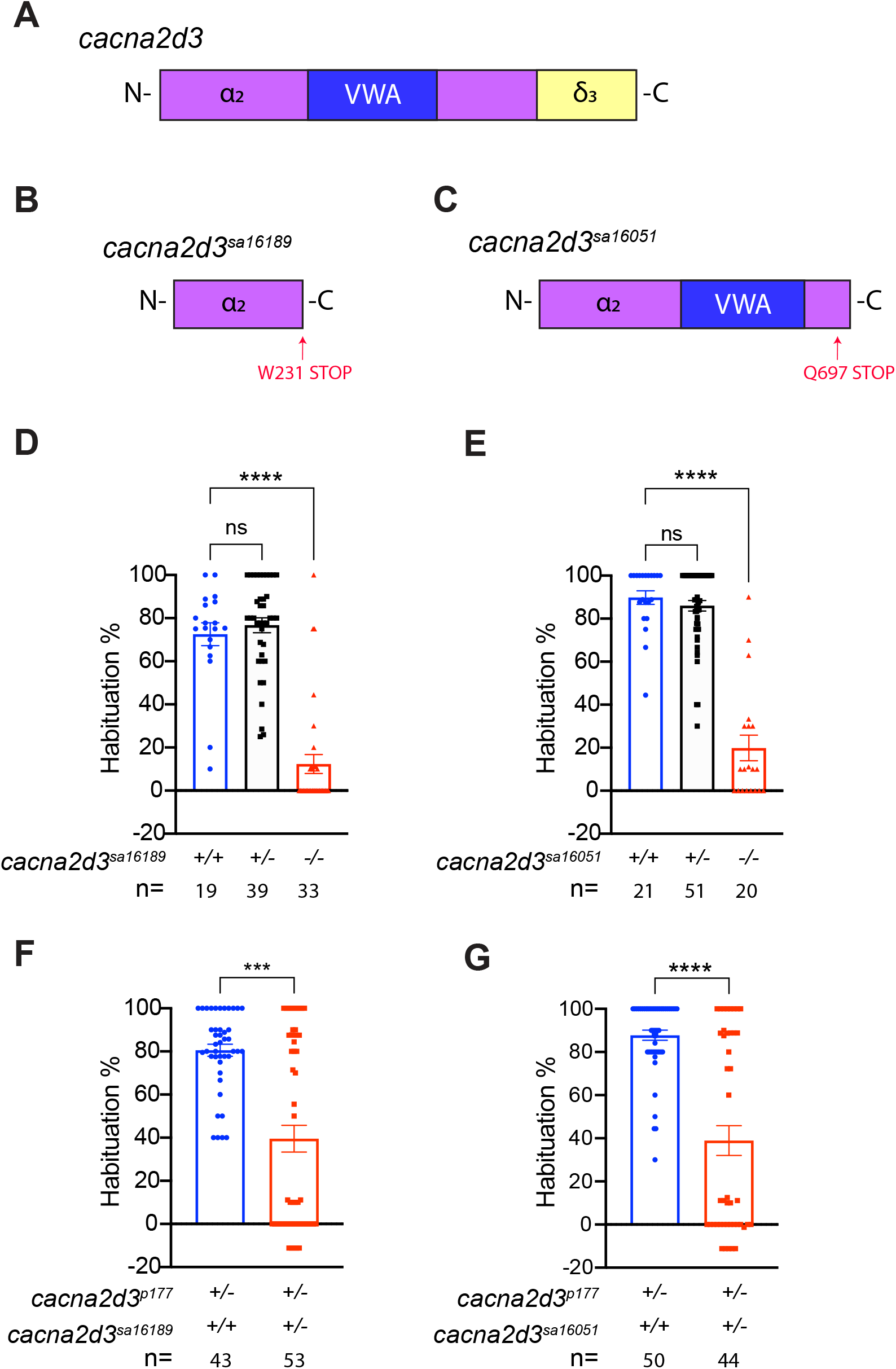
*cacna2d3* mutants and trans-heterozygotes exhibit reduced ASR habituation. (A-C) Schematics of the wildtype *cacna2d3* allele (A), the *cacna2d3*^*sa16189*^ mutant allele (B), and the *cacna2d3*^*sa16051*^ mutant allele (C). (D) Mean acoustic startle habituation percentage from an incross of *cacna2d3*^*sa16189*^*/+* heterozygotes. Wildtype larvae are shown in blue, *cacna2d3*^*sa16189*^*/+* heterozygotes are shown in black, and homozygous *cacna2d3*^*sa16189*^ mutants are shown in red. Number of larvae tested shown below each bar. (E) Mean acoustic startle habituation percentage from an incross of *cacna2d3*^*sa16051*^*/+* heterozygotes. Wildtype larvae are shown in blue, *cacna2d3*^*sa16051*^*/+* heterozygotes are shown in black, and homozygous *cacna2d3*^*sa16051*^ mutants are shown in red. (D-E) ****p<0.0001, one-way ANOVA with Dunnett’s multiple comparison test. Error bars indicate SEM. (F) Mean acoustic startle habituation percentage from an outcross of *cacna2d3*^*p177*^ mutants with *cacna2d3*^*sa16189*^*/+* heterozygotes. *cacna2d3*^*p177*^*/*+ larvae are shown in blue and *cacna2d3*^*p177*^/*cacna2d3*^*sa16189*^ trans-heterozygous are shown in red. (G) Mean acoustic startle habituation percentage from an outcross of *cacna2d3*^*p177*^ mutants with *cacna2d3*^*sa16051*^*/+* heterozygotes. *cacna2d3*^*p177*^/+ larvae are shown in blue and *cacna2d3*^*p177*^/*cacna2d3*^*sa16051*^ trans-heterozygotes are shown in red. (F-G) ***p<0.001, ****p<0.0001, Mann-Whitney test versus wildtype. Error bars indicate SEM.

### cacna2d3 regulates habituation learning

We measured the ASR of *dory*^*p177*^ mutant larvae versus wildtype (TL) larvae using an automated behavioral platform that delivers acoustic stimuli of defined intensity, records behavioral responses with a high-speed camera, and tracks each animal’s movement to evaluate the initiation and kinematic performance of ASR behavior (Burgess and Granato, 2007; Wolman et al., 2011). Larvae were exposed to a series of 50 acoustic stimuli (Figure 1D). The first 10 stimuli (the “sensitivity” phase) were delivered at a subthreshold low-level intensity and spaced at 30-second intervals to assess startle sensitivity. The intensity of these subthreshold weak stimuli was chosen empirically to elicit SLCs ∼10% of the time in wildtype larvae during this phase. The next 10 stimuli (the “pre-habituation” phase) were delivered at a high-level intensity and spaced at non-habituating intervals of 30 seconds to determine baseline acoustic startle responsiveness. The intensity of these strong stimuli was set empirically to elicit SLCs ∼80% of the time in wildtype larvae during this phase. The following 30 stimuli (the “habituation” phase) were delivered at the same high-level intensity, but spaced only 3 seconds apart, which elicits short-term habituation (STH). Under these conditions, wildtype (TL) larvae show a rapid reduction in SLC startle frequency and stereotypically habituate by more than 80% (calculated as a fraction of the final response frequency over the initial response frequency, Figure 1E). In this assay, *dory*^*p177*^ homozygotes exhibited startle habituation of only 5.4% compared to the wildtype control larvae, which habituated by 83.3% (Figure 1F; wildtype controls n=68, *dory* ^*p177*^ homozygotes n=63).

Although WGS analysis and homozygosity mapping show the *cacna2d3* mutation to be strongly linked to the *dory* ^*p177*^ habituation defect, they do not prove a causal relationship. To address causality, we examined two additional *cacna2d3* mutant alleles obtained from the Zebrafish Mutation Project of the Sanger Center (Busch-Nentwich, 2013). *cacna2d3*^*sa16189*^ harbors a nonsense mutation at codon 231 of 1095, upstream of both the VWA functional domain and the *δ-3* subunit (Figure 2B). *cacna2d3*^*sa16051*^ harbors a nonsense mutation at codon 697, downstream of the VWA functional domain but upstream of the *δ-3* subunit (Figure 2C).

Wildtype and *cacna2d3*^*sa16189*^*/+* heterozygous siblings showed a rapid reduction in SLC startle response frequency and stereotypically habituated by 72.5% and 76.7%, respectively (Figure 2D). In contrast, *cacna2d3*^*sa16189*^ homozygous mutants exhibited weak startle habituation of 12.4% (Figure 2D; wildtype siblings n=19, *cacna2d3*^*sa16189*^*/+* n=39, *cacna2d3*^*sa16189*^ n=33). Similarly, wildtype and *cacna2d3*^*sa16051*^*/+* heterozygous siblings showed a rapid reduction in startle initiation and habituated by 89.8% and 86.0%, respectively, while *cacna2d3*^*sa16051*^ mutants showed weak habituation of 19.9% (Figure 2E; wildtype siblings n=21, *cacna2d3*^*sa16051*^*/+* n=51, *cacna2d3*^*sa16051*^ n=20). Habituation defects linked to three independent mutant alleles of *cacna2d3* demonstrate definitively that *cacna2d3* function is required for habituation learning.

To confirm that the habituation defect in *dory*^*p177*^, now renamed *cacna2d3*^*p177*^, is indeed caused by the splice-site mutation identified in *cacna2d3*, we tested trans-heterozygous larvae for habituation. *cacna2d3*^*p177*^*/cacna2d3*^*sa16189*^ trans-heterozygotes showed 39.5% habituation compared with the 80.5% habituation exhibited by siblings (Figure 2F; siblings n=43, *cacna2d3*^*p177*^*/cacna2d3*^*sa16189*^ n=53). Similarly, *cacna2d3*^*p177*^*/cacna2d3*^*sa16051*^ trans-heterozygotes showed 39.0% habituation compared to the 87.8% habituation exhibited by siblings (Figure 2G; siblings n=50, *cacna2d3*^*p177*^*/cacna2d3*^*sa16051*^ n=44). Together, these data indicate that the *cacna2d3*^*p177*^ habituation defect is caused by the unique T to C substitution within the splice donor sequence of intron 2-3 of *cacna2d3*.

### cacna2d3 regulates the innate acoustic startle threshold

In the course of habituation analyses, we noted unusual sensitivity of the mutant larvae during the sensitivity (low intensity stimulus) phase of the assay. Compared to the low response rate observed in wildtype (TL) larvae, *cacna2d3*^*p177*^ mutants exhibited a marked increase in SLC startle responsiveness to subthreshold low-level intensity stimuli (Figure 3A; 9.3% SLC response rate wildtype larvae n=68, compared to 52.4% in *cacna2d3*^*p177*^ homozygotes n=63). Similarly, *cacna2d3*^*sa16189*^ mutants showed 70% SLC startle responsiveness to subthreshold low-intensity stimuli, contrasted with the 16% SLC startle responsiveness of the wildtype and heterozygous siblings (Figure 3B; wildtype siblings n=19, *cacna2d3*^*sa16189*^*/+* n=40, *cacna2d3*^*sa16189*^ n=33). *cacna2d3*^*sa16051*^ mutants showed 60% SLC startle responsiveness compared to 11% of their wildtype and heterozygous siblings (Figure 3C; wildtype siblings n=23 *cacna2d3*^*sa16051*^*/+* n=51, *cacna2d3*^*sa16051*^ n=21. This increased SLC startle sensitivity was also observed in *cacna2d3*^*p177*^*/cacna2d3*^*sa16189*^ trans-heterozygotes, which showed 42.4% SLC startle responsiveness compared to 9.4% of their siblings (Figure 3D; siblings n=49, *cacna2d3*^*p177*^*/cacna2d3*^*sa16189*^ n=44) and in *cacna2d3*^*p177*^*/cacna2d3*^*sa16051*^ trans-heterozygotes, which showed 38.2% SLC startle responsiveness compared to 4.8% of siblings (Figure 3E; siblings n=43, *cacna2d3*^*p177*^*/cacna2d3*^*sa16051*^ n=53). SLC startle sensitivity defects in trans-heterozygotes were consistently less pronounced than in homozygous larvae, possibly due to differences in genetic backgrounds between the strains in which these alleles were maintained. Collectively, these results are consistent with the notion that loss of *cacna2d3* leads to a lower threshold of the startle response.

**Figure 3.**
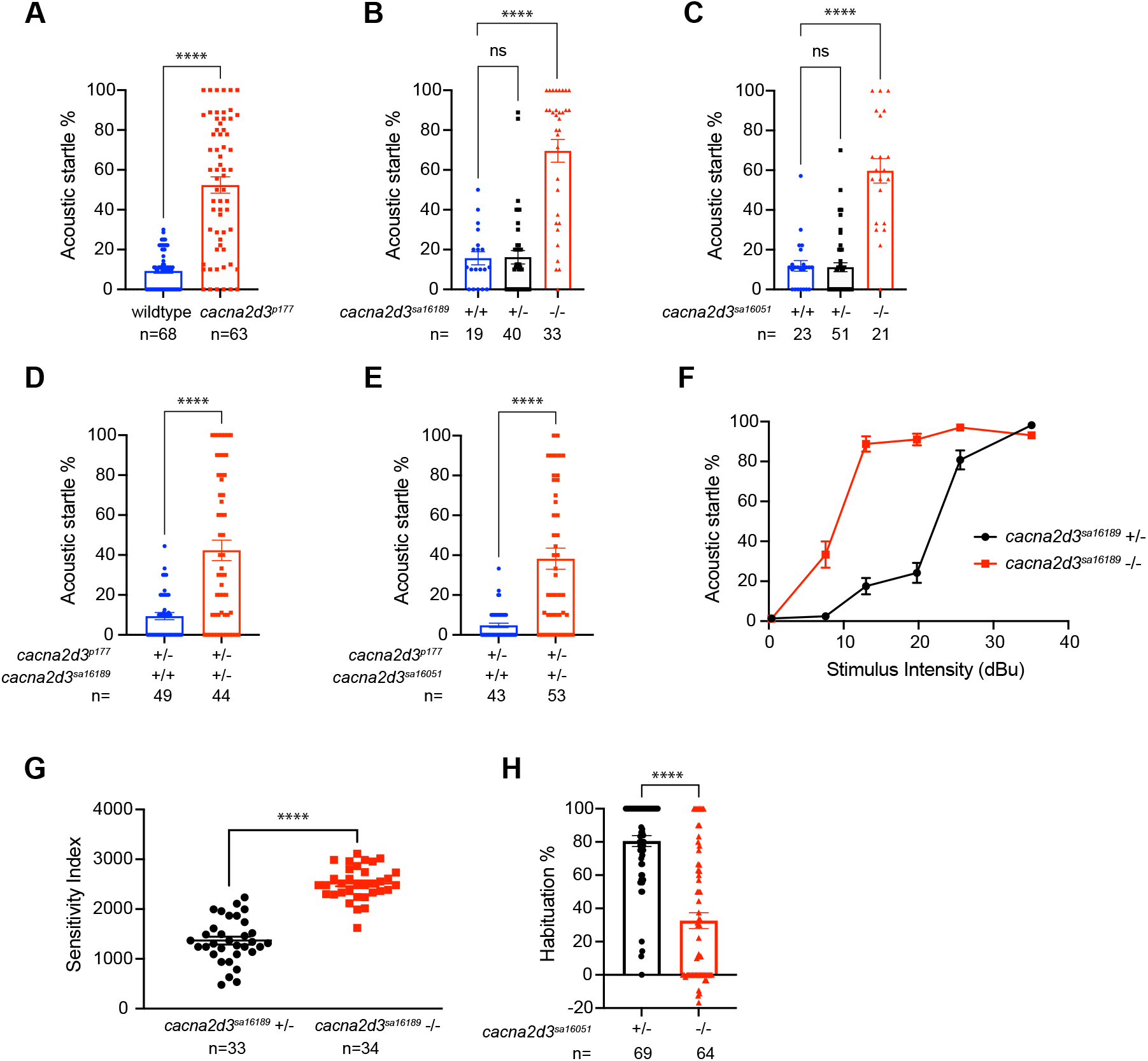
The startle threshold is reduced in *cacna2d3* mutants. (A-E) Acoustic startle (SLC) responsiveness of *cacna2d3* mutants to the 10, low-level acoustic stimuli presented during the “sensitivity” phase of the habituation assay. Startle response of (A) wildtype (shown in blue) and *dory*^*p177*^ mutants (shown in red). Startle response of (B) *cacna2d3*^*sa16189*^ wildtype (shown in blue), heterozygous (shown in black) and mutant larvae (shown in red). Startle response of (C) *cacna2d3*^*sa16051*^ wildtype (shown in blue), heterozygous (shown in black) and mutant larvae (shown in red). (B, C) ****p<0.0001, one-way ANOVA with Dunnett’s multiple comparison test. Number of larvae analyzed is shown below each bar. Error bars indicate SEM. Startle response of (D) *dory*^*p177*^*/+* sibling larvae (shown in blue) and *cacna2d3*^*p177*^/*cacna2d3*^*sa16189*^ trans-heterozygous larvae (shown in red). Startle response of *dory*^*p177*^*/+*; siblings (shown in blue) and *cacna2d3*^*p177*^/*cacna2d3*^*sa16051*^ trans-heterozygous larvae (shown in red). (A, D, E) ****p<0.0001, unpaired t-test with Welch’s correction versus wildtype. Number of larvae analyzed is shown below each bar. Error bars indicate SEM. (F) Startle frequency for 30 trials at 6 stimulus intensities. *cacna2d3*^*sa16189*^*/+* heterozygotes are in black and *cacna2d3*^*sa16189*^ mutants are in red. (G) Mean startle sensitivity indices. ****p<0.0001, unpaired t-test. Number of larvae analyzed is shown below dot plot. Error bars indicate SEM. (H) Mean acoustic startle habituation percentage from a cross of *cacna2d3*^*sa16051*^*/+* heterozygotes and *cacna2d3*^*sa16051*^ mutants with a lowered acoustic intensity. *cacna2d3*^*sa16051*^*/+* heterozygotes are shown in black, and homozygous *cacna2d3*^*sa16051*^ mutants are shown in red. ****p<0.0001, unpaired t-test with Welch’s correction versus heterozygotes. Error bars indicate SEM. Number of larvae analyzed is shown below each bar.

To test this hypothesis further, we subjected *cacna2d3*^*sa16189*^ mutants and their heterozygous siblings to varying intensities of acoustic stimuli at non-habituating intervals and measured SLC startle responsiveness. This analysis revealed that intensities as low as 7.6dB cause a significant increase in SLC startle responsiveness of 33.4% in *cacna2d3*^*sa16189*^ mutants, compared to 2.4% in heterozygous siblings (Figure 3F; *cacna2d3*^*sa16189*^*/+* n=33, *cacna2d3*^*sa16189*^ n=33). At 13dB, the SLC startle responsiveness in *cacna2d3*^*sa16189*^ mutants reaches 88.8%, a level that their heterozygous siblings do not reach until 25.6dB. To quantify the severity of this hypersensitivity phenotype, we calculated the startle sensitivity index by plotting the startle frequency of each larva across the 30-stimulus assay and measuring the area under the resulting curves. *cacna2d3*^*sa16189*^ homozygous mutants exhibited significant hypersensitivity compared to their heterozygous siblings (Figure 3G; *cacna2d3*^*sa16189*^*/+* n=33, *cacna2d3*^*sa16189*^ n=34). These data show that, in addition to its role in habituation, *cacna2d3* is required for establishing or maintaining acoustic startle thresholds.

To examine the relationship between the habituation defect in *cacna2d3* mutants and their behavioral hypersensitivity, we asked if their ability to habituate could be restored by lowering the intensity of habituating stimuli relative to the original habituation assay (Figure 1D). For this experiment, we used acoustic stimuli empirically determined to elicit SLCs ∼25% of the time in wildtype larvae. When subjected to the modified regime with these intermediate reduced-intensity stimuli, *cacna2d3*^*sa16051*^ mutants and their heterozygous siblings still showed significant learning impairments, with *cacna2d3*^*sa16051*^ mutants exhibiting 32.6% habituation compared to 80.5% in heterozygous siblings (Figure 3H; *cacna2d3*^*sa16051*^*/+* n=69, *cacna2d3*^*sa16051*^ n=64).

### cacna2d3 controls latency of the acoustic startle response

To investigate the motor function of mutant larvae, we assessed their SLC kinematics. The SLC maneuver executed in response to high intensity acoustic stimuli is defined by kinematic parameters within a well-defined range of values. These parameters include C-turn initiation latency and C-turn duration (Burgess and Granato, 2007). *cacna2d3*^*p177*^ mutants showed a marked reduction in C-turn latency compared to wildtype (TL) larvae (Figure 4A) but no difference in C-turn duration (Figure 4B). SLC kinematic analysis of the *cacna2d3*^*sa16189*^ and *cacna2d3*^*sa16051*^ mutants revealed that both mutants also showed a significant reduction in C-turn latency (Figures 4C, 4E, respectively), but no change in C-turn duration (Figures 4D, 4F, respectively). Lastly, in order to confirm a causative role for *cacna2d3* in regulating C-turn latency, but not C-turn duration, we evaluated the startle kinematics of the trans-heterozygotes in the complementation analysis, which revealed that *cacna2d3*^*p177*^*/ cacna2d3*^*sa16189*^ *larvae* exhibited reduced C-turn latency (Figure 4G) and no change in C-turn duration (Figure 4H).

**Figure 4.**
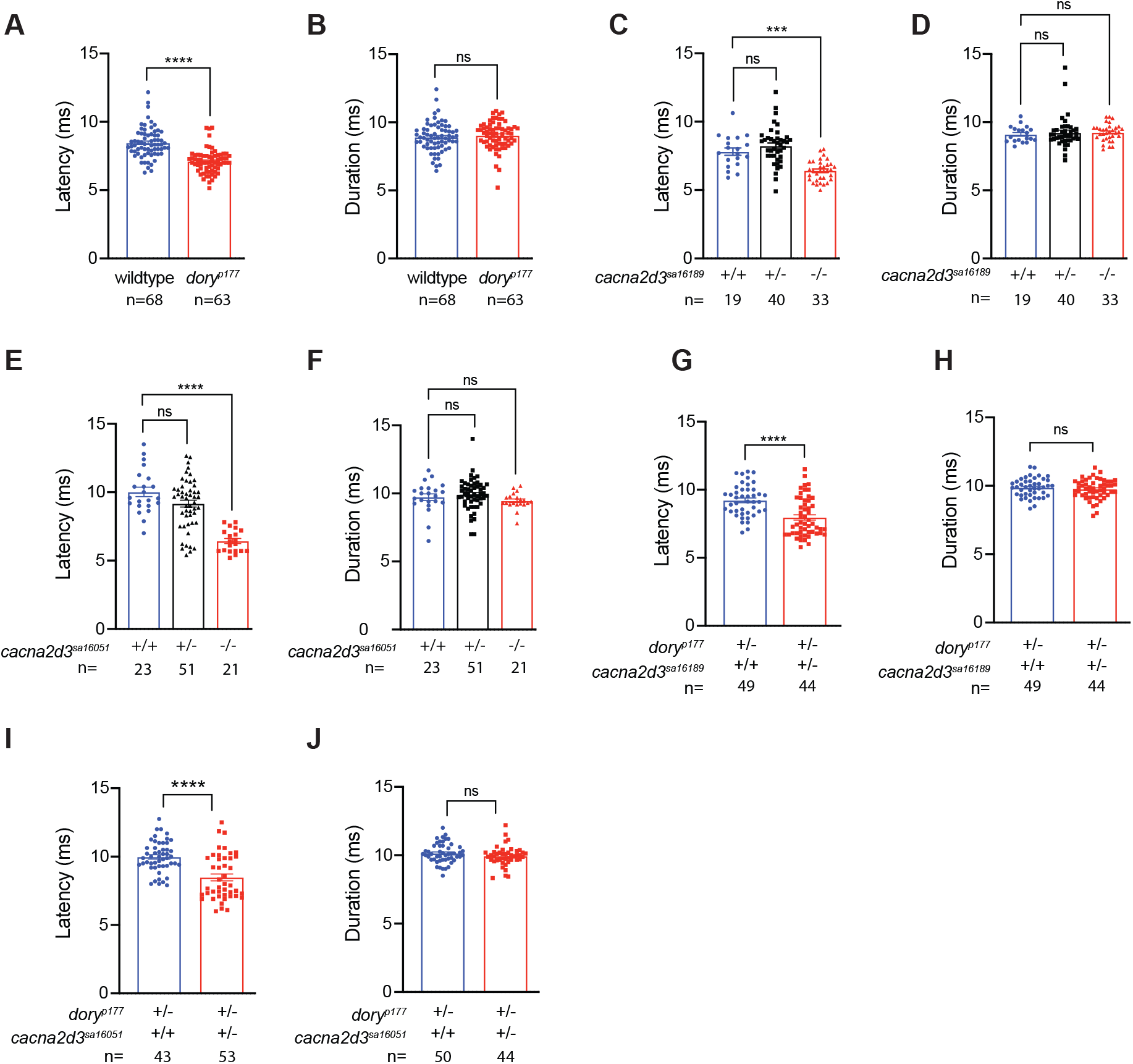
*cacna2d3* mutants and trans-heterozygotes exhibit impaired startle latency. Startle kinematic analysis of C-turn latency and C-turn duration. (A) Latency and (B) duration analysis and of wildtype (shown in blue) and *cacna2d3*^*p177*^ mutants (shown in red). (A-B) ns, ****p<0.0001, Mann-Whitney test versus wildtype. (C) Latency and (D) duration analysis of *cacna2d3*^*sa16189*^ wildtype (shown in blue), heterozygous (shown in black) and mutant larvae (shown in red). (E) Latency and (F) duration analysis of *cacna2d3*^*sa16051*^ wildtype (shown in blue), heterozygous (shown in black) and mutant larvae (shown in red). (C, E) ***p<0.001, ****p<0.0001, one-way ANOVA with Dunnett’s multiple comparison test versus wildtype. (D, F) ns, Kruskall-Wallis test with Dunn’s multiple comparisons test versus wildtype. (G) Latency and (H) duration analysis of *cacna2d3*^*p177*^*/+* sibling larvae (shown in blue) and *cacna2d3*^*p177*^/*cacna2d3*^*sa16189*^ trans-heterozygous larvae (shown in red). (I) Latency and (J) duration analysis of *cacna2d3*^*p177*^*/+* siblings (shown in blue) and *cacna2d3*^*p177*^/*cacna2d3*^*sa16051*^ trans-heterozygous larvae (shown in red). (G-I) ns, ****p<0.0001, unpaired t-test with Welch’s correction versus *cacna2d3*^*p177*^ +/-siblings. (J) ns, Mann-Whitney test versus *cacna2d3*^*p177*^*/+* siblings. Number of larvae analyzed is shown below each bar. Error bars indicate SEM.

Similarly, *cacna2d3*^*p177*^*/cacna2d3*^*sa16051*^ trans-heterozygotes showed reduced C-turn latency (Figure 4I) with no change in C-turn duration (Figure 4J). The decreased latency of the SLC is likely due to decreased startle threshold (Marsden et al., 2018), which leads them to initiate the escape maneuver more rapidly than their wildtype and heterozygous siblings. Despite these differences, these kinematic parameters are still within the range previously used to define SLC responses (Burgess and Granato, 2007) and are consistent with a normal motor function controlling escape responses in *cacna2d3* mutants.

## Discussion

Habituation is a fundamental form of learning that is conserved across species (Groves and Thompson, 1970; Rankin et al., 2009; Thompson and Spencer, 1966). It is defined as a learning process in which an organism’s responsiveness to a given stimulus progressively declines with repeated exposure. In humans, aberrant habituation is a hallmark of many behavioral disorders that exhibit cognitive dysfunction, including ASDs (Ornitz et al., 1993; Perry et al., 2007; Takahashi et al., 2016), schizophrenia (Braff and Geyer, 1990), and ADHD (Jansiewicz et al., 2004). The etiologies of these disorders are under intense scrutiny, and there is a critical need to identify genes important for habituation as candidate therapeutic targets for these disorders. Whole genome sequence analysis of the *dory/cacna2d3*^*p177*^ mutant line, obtained in a forward-genetic screen for habituation mutants (Wolman et al., 2015), led us to uncover a previously unknown role for *cacna2d3* in habituation to acoustic stimuli in vertebrates, and in establishing or maintaining a baseline innate startle threshold, thus altering sensitivity to acoustic stimuli.

*cacna2d3* encodes the calcium voltage-gated channel auxiliary subunit alpha-2/delta-3. Alpha-2/delta subunits function at the presynaptic terminal by strengthening the coupling between calcium influx and neurotransmitter release (Hoppa et al., 2012). The α2 and δ3 subunits are generated through proteolytic cleavage of a precursor protein encoded by *cacna2d3* (Dolphin, 2013). In the ER, a GPI anchor is added to the δ3 protein, which anchors the subunit to the cell membrane (Davies et al., 2010). Cacna2d3 is highly conserved across vertebrates, including zebrafish (Supplemental Figure 1), particularly in the Von Willebrand Factor A (VWA) domain that is involved in protein-protein interactions via a metal ion adhesion (MIDAS) motif (Dolphin, 2013) and is important for trafficking of VGCCs (Canti et al., 2005).

In addition to impaired habituation, we have documented enhanced sensitivity to acoustic stimuli in *cacna2d3* mutants (Figures 3A-E). This is evidenced by an increased sensitivity index, suggesting their innate startle threshold is reduced (Figures 3G). The innate threshold for the startle response is an important mechanism for regulating threat evasion (Marsden et al., 2018); reduction of this threshold and the subsequent hypersensitivity to acoustic stimuli has been strongly linked to ASD (Takahashi et al., 2016) and anxiety in humans (Bakker et al., 2009). The intensity of a stimulus determines how rapidly the habituation will occur, with weaker stimuli resulting in a more pronounced reduction in responsiveness than robust stimuli (Rankin et al., 2009). It will be interesting to investigate the potential causal relationship between increased sensitivity and decreased habituation in future studies.

Mutations in the *C. elegans* ortholog of *cacna2d3, unc-36*, have been linked with impaired habituation to tactile stimuli, as well as increased tactile sensitivity (McDiarmid et al., 2020). In combination with our findings that *cacna2d3* mutant zebrafish show impaired habituation to acoustic stimuli (Figures 1E, 2B and 2F), these data suggest a strong functional conservation of *cacna2d3* in habituation learning. In contrast to *cacna2d3* mutant zebrafish, mice with *CACNA2D3* dysfunction exhibit reduced ASR when presented with acoustic stimuli and have a higher startle threshold (Pirone et al., 2014). This reduction in ASR is linked to reduced levels of presynaptic calcium channels and smaller synaptic boutons in auditory nerve fibers.

Remarkably, *CACNA2D3* mutant mice show a marked increase in tactile startle responsiveness when presented with stimuli elicited by air puffs. The increased startle responsiveness to tactile stimuli in mice and *C. elegans* is of particular interest, as tactile hypersensitivity has been linked to anxiety, autism, and migraines (McDiarmid et al., 2017; Orefice et al., 2016). While zebrafish detect acoustic stimuli via the hair cells in their otic vesicles, as do other animals, the zebrafish lateral line also contains hair cells that detect vibrations in the water and encode them as mechanosensory stimuli (Bleckmann and Zelick, 2009). It may be that, similar to the tactile hypersensitivity observed in *C. elegans* (McDiarmid et al., 2020) and mice (Pirone et al., 2014), hypersensitivity observed in *cacna2d3* mutant zebrafish is due to a combination of acoustic-and mechanosensory-driven aberrant startle responses. Accessibility of the lateral line hair cells to pharmacological manipulation and ablation offers an effective strategy to test this hypothesis in the future (Alassaf et al., 2019).

Together, our findings identify essential functions for zebrafish *cacna2d3* in acoustic startle sensitivity and habituation. The high degree of sequence conservation suggests strong conservation of Cacna2d3 protein functions from zebrafish to human and supports the value of *cacna2d3* mutant zebrafish as a clinically relevant model for elucidating the underlying mechanisms of sensory filtering impairments associated with prevalent neurodevelopmental disorders.

## Methods

### Generation and Maintenance of Zebrafish

Zebrafish (Danio rerio) were maintained according to established methods (Westerfield, 1993). All experimental protocols using zebrafish were approved by the University of Wisconsin Animal Care and Use Committee and carried out in accordance with the institutional animal care protocols. Embryos were generated from natural matings of adult Tüpel long fin (TL), and adults carrying the *dory/cacna2d3*^*p177*^, *cacna2d3*^*sa16051*^, and *cacna2d3*^*sa16189*^ alleles, respectively.

Embryos were raised in E3 media at 28°C on a 14 hr/10 hr light/dark cycle through 5 dpf as previously described (Kimmel et al., 1995, Gyda et al., 2012). 5 dpf larvae were analyzed for behavior in E3.

### Genotyping

To genotype larvae, we developed dCAPS assays using the dCAPS program (http://helix.wustl.edu/dcaps/dcaps.html) to design appropriate primers (Neff et al., 2002) for the *dory*^*p177*^ and *cacna2d3*^*sa16051*^ alleles. Primers for the *cacna2d3*^*sa16189*^ allele were designed using Primer3 (Untergasser et al., 2007). Primer sequences, PCR conditions, and restriction endonucleases used for digestion are outlined in **Table 1**. All genotyping was performed after behavioral experiments. *cacna2d3*^*sa16189*^ larvae used in sensitivity assay were genotyped using the KASP method with proprietary primer sequences (LGC Genomics).

**Table 1.**
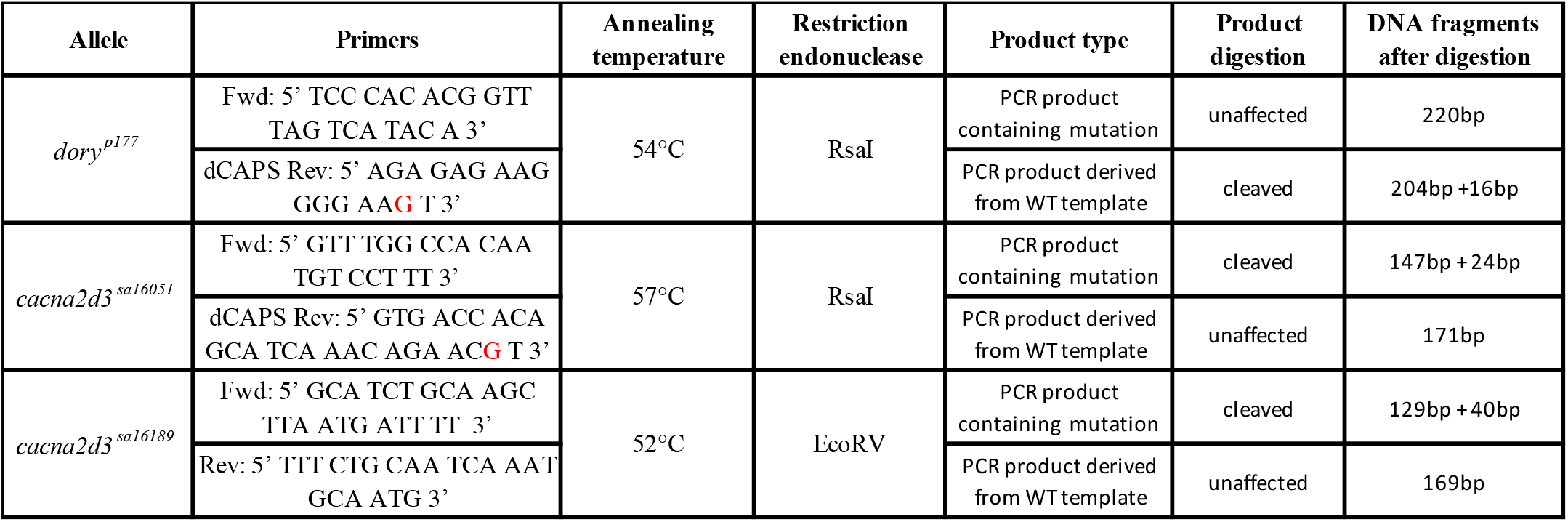
Genotyping primers and genotyping conditions. Genotyping primers, annealing temperatures, restriction endonuclease used for digestion, and expected band fragment sizes are listed. A mismatch (marked in red) has been introduced into the reverse primer for the *dory*^*p177*^ allele that creates an RsaI restriction enzyme site in the amplified product from the wildtype DNA template. Similarly, a mismatch (marked in red) has been introduced into the reverse primer for the *cacna2d3*^*sa16051*^ allele that creates an RsaI restriction enzyme site in the amplified product from the mutant DNA template.

### Behavioral Analyses

On the day of habituation behavioral testing and acoustic sensitivity analysis, larvae were held in 60mm-wide Petri dishes with 25 larvae in 10mL E3, kept on a white light box for at least 30 minutes, and then transferred to a 6×6 grid. Startle behavior was elicited using an automated behavioral platform in which the intensity and timing of acoustic stimuli could be controlled (Burgess and Granato, 2007; Wolman et al., 2011). Startle responses were elicited with a minishaker (Brüel & Kjær, Model 4810). For the habituation assays, the acoustic stimuli were of 3 millisecond duration, with 1000 Hz waveforms, at either low-level, subthreshold intensity identified empirically to elicit SLCs ∼10% of the time in wildtype larvae or above threshold, high intensity identified empirically to elicit SLCs ∼80% of the time in wildtype larvae. For the experiment examining the relationship between the habituation defect in *cacna2d3* mutants and their behavioral hypersensitivity, we used an intermediate level acoustic stimulus selected empirically to elicit SLCs ∼25% of the time in wildtype larvae.

The habituation assay consists of multiple phases, each designed to assess different parameters of short-term acoustic startle habituation. To evaluate acoustic sensitivity, low-level “subthreshold” intensity stimuli were presented at a 30 second interstimulus interval (ISI) (stimuli 1-10), eliciting ∼10% startle responses in wildtype larvae. To evaluate short-term startle habituation, high-intensity stimuli were presented at 30 second ISI during the “pre-habituation” phase (stimuli 11-20) and at 3 second ISI during the “habituation” phase (stimuli 21-50). The degree to which larvae habituate was calculated by comparing the average frequency of startle responsiveness of an individual during the pre-habituation and the last 10 stimuli of the habituation phases (stimuli 41-50) (Wolman et al., 2011).

Startle responses were captured at 1000 frames per second with a MotionPro Y4 video camera (Integrated Design Tools) with a 50 mm macro lens (Sigma Corporation of America) at 512 × 512 pixel resolution. We used FLOTE to analyze startle responses in an experimenter-independent, automated manner (Burgess and Granato, 2007). FLOTE tracks the position of individual larvae frame by frame and characterizes locomotor maneuvers (e.g. C-bend, routine turn, swim, etc) according to predefined kinematic parameters that distinguish these maneuvers. We used a custom R-script to run analysis on behavior data generated by FLOTE to calculate habituation and analyze kinematic data, which allowed for an additional level of experimenter-independent, automated analysis. For startle behavior, we report data representing the short-latency C-bend (SLC) startle response. When testing individual larvae for habituation, animals that exhibited a startle response of <40% to acoustic stimuli during the “pre-habituation” phase were classed as non-responders and excluded from analysis. For kinematic data, we report the SLC response of larvae during the 10 high-intensity stimuli given during the “pre-habituation” phase of the habituation assay.

For the generation of sensitivity index calculations, *cacna2d3*^*sa16189*^ homozygous mutants were crossed with *cacna2d3*^*sa16189*^*/+* heterozygous carriers. Larvae were tested for acoustic behavioral sensitivity at 5dpf and analyzed using FLOTE software as described previously (Burgess and Granato, 2007; Marsden et al., 2018). Briefly, larvae were presented with a total of 30 acoustic stimuli: 5 trials of 6 different stimulus intensities at the following decibel levels: 0.3 dB, 7.6 dB, 13 dB, 19.8 dB, 25.6 dB, and 35 dB. Each stimulus was separated with a 40-second ISI. Percent startle for each larva was recorded at each stimulus intensity. Sensitivity index was calculated for each larva by calculating the area under the curve of percent startle vs. stimulus intensity using Prism (GraphPad). Stimulus intensities were calibrated using a PCB Piezotronics accelerometer (#355B04) and signal conditioner (#482A21). Voltage outputs were converted to dBu using the formula dBu = 20* log (V/0.775).

### Whole Genome Sequencing

Positional cloning was performed as previously described (Wolman et al., 2015). A pool of 64 behaviorally identified *dory* mutant larvae was collected, genomic DNA (gDNA) was extracted, and gDNA libraries were prepared. gDNA was sequenced with 100-bp paired-end reads on the Illumina HiSeq 2000 platform, and homozygosity analysis was done using 463,379 SNP markers identified by sequencing gDNA from ENU-mutagenized TL and WIK males as described previously (Wolman et al., 2015).

### Statistics

All graph generation and statistical analyses, including calculation of means and SEM, were performed using Graphpad Prism (www.graphpad.com). D’Agostino and Pearson normality test was used to test whether data were normally distributed. If data were normally distributed, significance was assessed using t-tests with Welch’s correction or ANOVA with Dunnet’s multiple comparisons test. If data were not normally distributed, Mann-Whitney test or Kruskall-Wallis test with Dunn’s multiple comparisons test was used.

## Acknowledgements

We thank the members of the Grinblat lab for technical support and the members of the labs of Dr. Mary Halloran, Dr. Katie Drerup, and Dr. David Ehrlich for zebrafish line maintenance. This work was funded by grants from the University of Wisconsin-Madison (Science and Medicine Graduate Research fellowship to N.J.S., Department of Integrative Biology funding to A.H.M. and N.J.S, and the Office of the Vice Chancellor for Research and Graduate Education/Wisconsin Alumni Research Foundation to N.J.S. and Y. G), and by the National Science Foundation Graduate Research Fellowship Program under Grant Numbers DGE-1256259 and DGE-1747503 to N.J.S., NIH K99NS111736 to J.C.N., NIDCD 5T32DC016903 to E.A.O., and NIH R01NS118921 to M.G. Any opinions, findings, and conclusions or recommendations expressed in this material are those of the authors and do not necessarily reflect the views of the National Science Foundation.

## Author contributions

N.J.S., J.C.N., E.A.O, and Y.G. designed the experiments. N.J.S., E.A.O., D.K.H., and Z.S. performed the behavioral experiments. N.J.S. and E.A.O. analyzed the behavioral experiments with assistance from A.H.M. J.C.N. performed whole genome sequencing analysis and homozygosity mapping. A.H.M. developed the custom R-Script used for behavioral analysis. N.J.S. and Y.G. wrote the manuscript with input from all authors. Y.G. and M.G. directed the project.

## Figures and Tables

**Supplemental Figure 1**.

The alignment between human CACNA2D3 and zebrafish Cacna2d3 protein sequence was generated using the local alignment algorithm (Smith-Waterman) in SnapGene software. Human CANCA2D3 and zebrafish Cacna2d3 proteins share 76.5% amino acid identity and 87.6% similarity along the length of the protein. In the Von Willebrand Factor A (VWA) functional domain, the proteins share 89.4% amino acid identity and 93.4% similarity. Human CACNA2D3 is designated by row 1 and zebrafish Cacna2d3 is designated by row 2. VWA domain designated by blue box. | = identical amino acid;: = similar amino acid;. = not similar amino acid

