## Supplemental Figure 1 for "*cacna2d3*, a voltage-gated calcium channel subunit, functions in vertebrate habituation learning and the startle sensitivity threshold"

Consensus
1. ENSP00000419101_Homo Sapiens
2. ENSDARP00000001281_Danio rerio

Legend: | Identical  : Similar  · Not Similar  VWA Domain

```
1   11    SRGASALLAAALLYAALG--DVVRSEQQIPLSVVKLWASAFGGEIKSIAAKYSGSQLLQK   68
2   4     NRRNPALLWCVMLSACWTPLEVRGSQPEIPLSVVKLWASAFGGEIKSIASKYSGSQLLQK   63

1   69    KYKEYEKDVAIEEIDGLQLVKKLAKNMEEMFHKKSEAVRRLVEAAEEAHLKHEFDADLQY   128
2   64    KYKEYEKSVRIEEIDGAKLVKNLAQNMEEMFRKKAEATRRLVEAAEEAHLQHEENPDLQY   123

1   129   EYFNAVLINERDKDGNFLELGKEFILAPNDHFNNLPVNISLSDVQVPTNMYNKDPAIVNG   188
2   124   EYFNAVLINEVDEDGNSVELGGEFLLEPNDHFNNLSVNLSLSVVQVPTNMYNKDPDIVNG   183

1   189   VYWSESLNKVFVDNFDRDPSLIWQYFGSAKGFFRQYPGIKWEPDENGVIAFDCRNRKWYI   248
2   184   VYWSEALNKVFVDNFRRDPTLIWQYFGSAKGFFRQYPGVKWYPDEHGVIAFDCRNRKWYI   243

1   249   QAATSPKDVVILVDVSGSMKGLRLTIAKQTVSSILDTLGDDDFFNIIAYNEELHYVEPCL   308
2   244   QAATSPKDVVILVDVSGSMKGLRLTIARQTVASILDTLGDDDFFNIIAYNQEIHYVEPCL   303

1   309   NGTLVQADRTNKEHFREHLDKLFAKGIGMLDIALNEAFNILSDFNHTGQGSICSQAIMLI   368
2   304   NGTLVQADSTNKDHFKEHLDKLFAKGIGLLGNALSEAFTILNEINQTGRGSSCSQAIMLI   363

1   369   TDGAVDTYDTIFAKYNWPDRKVRIFTYLIGREAAFADNLKWMACANKGFFTQISTLADVQ   428
2   364   TDGATEMYDDVFAKYNWPERKVRIFPYLIGRESAFADNLKWMACANKGYFSQISTLADVQ   423

1   429   ENVMEYLHVLSRPKVIDQEHDVVWTEAYIDSTLPQAQKLTDDQGPVLMTTVAMPVFSKQN   488
2   424   ENVMRYLHVMSRPKVIDHEHDTVWTEAYVDSALAQAHKLGENKGPNPMTTVAMPVFSTKN   483

1   489   ETRSKGILLGVVGTDVPVKELLKTIPKYKLGIHGYAFAITNNGYILTHPELRLLYEEGKK   548
2   484   ETKNQGILLGVAGTDIPIQELMKVIPKHKLGIHGYAFAITSNGYLLLHPDLRPLYQEGQK   543

1   549   RRKPNYSSVDLSEVEWEDRDDVLRNAMVNRKTGKFSMEVKKTVDKGKRVLVMTNDYYYTD   608
2   544   RKKPSYSSVDLSEVEWEDTEDVLRTAMVNRKTGTFSMEVKRTVDKGKRVLTMHNDYYYTD   603

1   609   IKGTPFSLGVALSRGHGKYFFRGNVTIEEGLHDLEHPDVSLADEWSYCNTDLHPEHRHLS   668
2   604   IKGTPFSVGVALSRGHGKYFFKGNVSLEAGLHDLEQPDVALADEWTYCTTEEHHEHRHLT   663

1   669   QLEAIKLYLKGKEPLLQCDKELIQEVLFDAVVSAPIEAYWTSLALNKSENSDKGVEVAFL   728
2   664   QVQAIKIYMTSRKPHFKCDRELIQQVLFDAVVTAPVEAYWTGLALNKSENSDKGVEIAFL   723

1   729   GTRTGLSRINLFVGAEQLTNQDFLKAGDKENIFNADHFPLWYRRAAEQIPGSFVYSIPFS   788
2   724   GTRTGLSRTNLFVVPEQLTNRDFLTAEDKEGVFNADHFPLWYKRAAEQVPGTFVYSIPFS   783

1   789   TGPVNKSNVVTASTSIQLLDERKSPVVAAVGIQMKLEFFQRKFWTASRQCASLDGKCSIS   848
2   784   SGGENKS-VVLASTAIQLLDDRKSPIVAAVGIQMKLEFFQRKFWTASRQCNALDGKCTIS   842

1   849   CDDETVNCYLIDNNGFILVSEDYTQTGDFFGEIEGAVMNKLLTMGSFKRITLYDYQAMCR   908
2   843   CDNEDIKCYLIDNNGFILVSEDTSQTNLFFGKVEGAVMNKLLLMGSFKKINLYDYQALCK   902

1   909   ANKESSDGAHGLLDPYNAFLSAVKWIMTELVLFLVEFNLCSWWHSDMTAKAQKLKQT-LE   967
2   903   EYAGSSDSARTLLDPF----TTVKWLLTELVIFLLEFNLYSWWNCDSTVKAQRSQRTMMV   958

1   968   PCDTEYPAFVSERTIKETTGNIACEDCSKSFVIQQIPSSNLFMVVVDSSCLCESVAPITM   1027
2   959   PCDTEYPAFVSERTIKETTGNIDCNGCIRTFVIQQIPSSNLFMVVVENKCDCGSALPVTM   1018

1   1028  APIEIRYNESLKCERLKAQKIRRRPESCHGFHPEENARECGGAPSLQAQTVLLLLPLLLM   1087
2   1019  EPIEIIYNESLKCDRLKFQKDRRRPQSCHPFHPEENAVECGSANALSFPLVAILIPALSL   1078

1   1088  LFSR   1091
2   1079  MISR   1082
```
